# A high-light tolerant alga from the desert is protected from oxidative stress by NPQ-independent responses

**DOI:** 10.1101/2024.06.10.598256

**Authors:** Guy Levin, Michael Yasmin, Oded Liran, Rawad Hanna, Oded Kleifeld, Guy Horev, Francis-André Wollman, Gadi Schuster, Wojciech J. Nawrocki

## Abstract

Non-photochemical quenching (NPQ) mechanisms are crucial for protecting photosynthesis from photoinhibition in plants, algae, and cyanobacteria, and their modulation is a long-standing goal for improving photosynthesis and crop yields. The current work demonstrates that *Chlorella ohadii*, a green micro-alga that thrives in the desert under high light intensities which are fatal to many photosynthetic organisms, does not perform nor require NPQ to protect photosynthesis under constant high light. Instead of dissipating excess energy, it minimizes its uptake by eliminating the photosynthetic antenna of photosystem II. In addition it accumulates antioxidants that neutralize harmful reactive oxygen species (ROS) and ramps up cyclic electron flow around PSI. These NPQ-independent responses proved efficient in preventing ROS accumulation and reducing oxidative damage to proteins in high-light-grown cells.

## Introduction

Light drives photosynthesis, but it also damages the photosynthetic apparatus in a number of ways. Particularly, illumination facilitates the generation of reactive oxygen species (ROS) which damage the molecular players in photosynthesis, thereby inhibiting the process – an effect known as photoinhibition (Ögren, 1988; Ögren and Sjöström, 1990). Under high light (HL) conditions, non-photochemical quenching (NPQ) mechanisms through which excess absorbed light energy is harmlessly dissipated as heat, become critical for photosynthesis protection (photoprotection)(Delosme, 1967; Wolff and Witt, 1969; Demmig-Adams et al., 1989; Demmig-Adams et al., 1990). The major NPQ component, q_E_, is located within the membrane-intrinsic major light-harvesting complex (LHCII) for photosystem II (PSII) and is activated by the acidification of the lumen compartment which results from photosynthetic electron transport. Typically, q_E_ is the strongest and the most rapidly triggered component of NPQ in the green lineage, undergoing activation in seconds, as recently reviewed by (Bassi and Dall’Osto, 2021). During periods of high photosynthetic activity, the thylakoid lumen is acidified due to proton translocation across the thylakoid membrane by the combined activity of linear and cyclic photosynthetic electron flows (LEF and CEF). PsbS and LhcSR in plants and algae, respectively act as proton sensors which, upon acidification of the lumen, induce q_E_ to dissipate excess energy as heat (Li et al., 2000; Li et al., 2004; Peers et al., 2009; Liguori et al., 2013). Simultaneously, the xanthophyll cycle is activated, and newly formed zeaxanthin interacting in the PSII-LHCII chlorophyll pool provides pathways for energy dissipation. (Demmig et al., 1987; Niyogi et al., 1998; Xu et al., 2015; Sacharz et al., 2017).

*Chlorella ohadii* is a green micro-alga that thrives in the desert, where it is exposed to extremely high light intensities (Treves et al., 2013; Levin et al., 2021). Accordingly, *C. ohadii* is highly tolerant to photoinhibition (Treves et al., 2016; Levin et al., 2023; Levin et al., 2024). The adaptation of *C. ohadii* to HL involves marked changes in thylakoid content of which the most pronounced is a decrease in LHCII, as well as the massive accumulation of carotenoids and several photoprotective proteins (Levin et al 2021, 2023). HL-grown *C. ohadii* does not undergo state transitions, a mechanism of antenna exchange between PSII and PSI (Bonaventura and Myers, 1969; Chow et al., 1981; Delosme et al., 1996), which also has been implicated in photoprotection (Allorent et al., 2013; Nawrocki et al., 2016). Moreover, *C. ohadii* also lack genes encoding LhcSR proteins nor does it accumulate the PsbS protein (Treves et al., 2016; Levin et al., 2023; Murik et al., 2024), suggesting the absence of q_E_. Interestingly, a carotenoid biosynthesis-related protein (CBR), which belongs to the LHC-like protein family with a structure similar to PsbS, accumulates to a great extent in HL-grown *C. ohadii* (Levy et al., 1993; Levin et al., 2023; Levin and Schuster, 2023). Structural modeling suggested that one of the proton-sensing glutamates of PsbS was conserved in CBR, raising the possibility of its role in lumen acidification sensing. Moreover, fractionation of the photosynthetic protein complexes by sucrose density gradient centrifugation showed that CBR from HL-grown cells co-localized with a detached fraction of LHCII and carotenoids, both of which are associated with NPQ and ROS scavenging (Levin et al., 2023). Given that LHCII is the major site of NPQ (Ruban et al., 1999; Elrad et al., 2002; Sacharz et al., 2017; Nicol and Croce, 2021), its elimination and the lack of LhcSR raise doubts as to the contribution of a q_E_ regulation which require to investigate further the mode of photoprotection in *C. ohadii*.

In this work, the contribution of the decrease in PSII antenna to HL acclimation, as well as the interplay between NPQ and CEF on ROS accumulation and oxidative damage in *C. ohadii* are investigated. We demonstrate that q_E_ is actually absent and replaced with an NPQ-independent photoprotection in HL-grown *C. ohadii* cells. This mechanism, rather than allowing energy dissipation, mainly relies limiting the absorption of light. It is concomitantly coupled to a dramatic enhancement of CEF around PSI and a vast accumulation of CBR, carotenoids, and other antioxidants, functionally reducing ROS accumulation. Thus, *C. ohadii,* adapted to grow at extremely high light intensity, employs both active and passive mechanisms of resistance to photoinhibition. The present work demonstrates that NPQ is not required to withstand excess illumination in HL-grown *C. ohadii* and that under conditions of prolonged exposure to HL intensities, minimizing the PSII-LHCII absorption cross-section and ROS production while maximizing antioxidant activity has been selected as the most successful strategy to reduce photoinhibition.

## Results

### HL-grown cells upregulate CBR and enzymatic antioxidant enzymes

To understand the remarkable resistance of *C. ohadii* to HL intensities, we grew it under low and high light conditions (LL: 50 µmol photons m^-2^ s^-1^; HL: 2000 µmol photons m^-2^ s^-1^), and then subjected the cells to label-free quantification mass spectrometry (LFQ-MS) to detect possible enzymatic antioxidants that participate in the detoxification of ROS and determine their effect on chloroplast protein oxidation. As was observed for isolated thylakoid membranes (Levin et al., 2023), the most prominent protein in HL-grown cells was CBR, with a 995-fold change compared to LL-grown cells. This was accompanied by a drastic loss in LHCII (Fig. 1A). Changes in LHCI abundance were much smaller than in LHCII, which suggests that PSII light-harvesting was more affected than PSI. Additionally, we identified an increase in 15 proteins associated with ROS detoxification or with energy dissipation via alternative electron transfer pathways, (Figs. 1A and 1B). Specifically, we detected two isoforms of superoxide dismutase (SOD), which catalyzes the detoxification of superoxide (O_2_^-^) by converting it to H_2_O_2,_, one of which showing a 1.8 higher accumulation in HL-grown cells than in LL-grown cells. Of glutathione peroxidase (GPX) and ascorbate peroxidase (APX), which can further detoxify H_2_O_2_ by catalyzing its conversion to water (H_2_O), only GPX was significantly accumulated in the HL-grown cells (3.3-fold change). Dehydroascorbate reductase (DHAR) and two isoforms of glutathione-s-transferase (GST), which participate in detoxifying the ascorbate-glutathione cycle, were also significantly accumulated in HL-grown cells (3.8-, 1.9- and 1.2-fold change, respectively). There were no statistically significant changes for catalase, which catalyzes the conversion of H_2_O_2_ to H_2_O in HL-grown cells. ω-3 fatty acid desaturase (ω3FAD), which catalyzes the generation of polyunsaturated fatty acids (PUFAs), underwent a 2.1-fold increased accumulation in HL-compared to LL-grown cells. PUFAs have been suggested to contribute to ROS detoxification by interacting with ROS during lipid peroxidation (Schmid-Siegert et al., 2016). Flavodiiron proteins (Flv1 and Flv2), which act as electron sinks possibly involved in PSII and/or PSI photoprotection (Chaux et al., 2015; Burlacot et al., 2022; Bag et al., 2023), were significantly accumulated in HL-grown cells (2.9- and 8.4-fold change, respectively), providing an additional route for energy dissipation. In contrast, PTOX, another putative electron sink (Nawrocki et al., 2015), was not differentially accumulated in HL-grown cells (0.9-fold change) (Figs. 1A and 1B).

**Figure 1.**
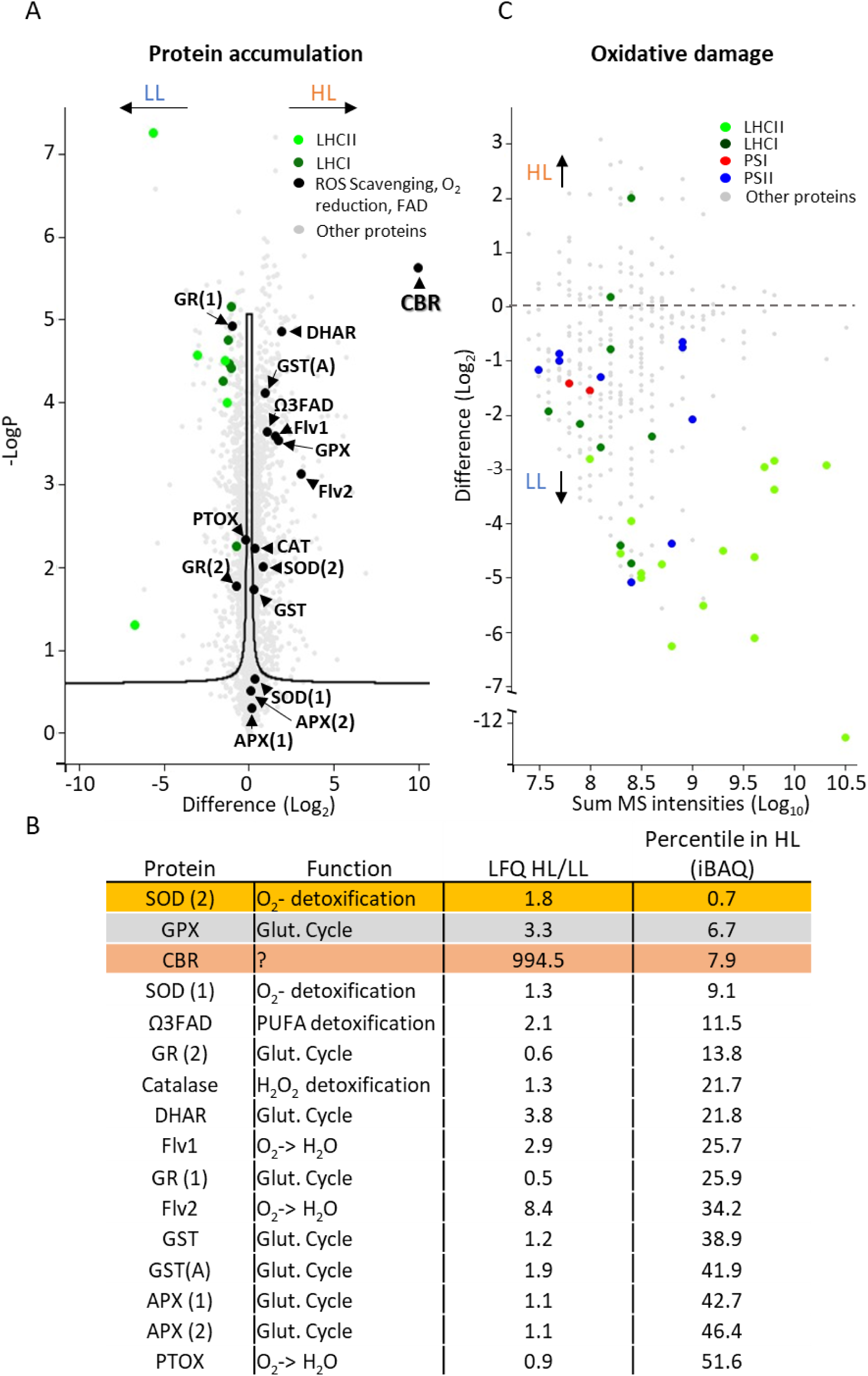
Adaptation to high light conditions in *Chlorella ohadii* as revealed by mass spectrometry. **A)** Label-free quantification (LFQ) Liquid chromatography-tandem mass spectrometry (LC-MS/MS) in low and high light (LL and HL)-grown cells reveals the accumulation of known enzymatic antioxidants and CBR in HL-grown cells. Note the substantial reduction in LHCII subunits in HL-grown *C. ohadii*. **B)** List of the detected antioxidants and other related enzymes. Values in the ‘LFQ HL/LL’ column represent the fold-change value of a given protein in HL over LL-grown cells. Values in the ‘Percentile in HL (iBAQ)’ column represent the abundance of the protein within HL-grown cells. A value of 10 means this protein is amongst the 10% most abundant proteins in these cells. **C)** Detected peptides with oxidized tryptophan as a mark for oxidative damage. Note the accumulation of almost all detected oxidized peptides in LL-grown cells. SOD – Superoxide dismutase. GPX – Glutathione peroxidase. CBR – Carotenoid biosynthesis related. Ω3FAD – omega 3 fatty acid desaturase. GR – Glutathione reductase. DHAR - Dehydroascorbate reductase. FLV – Flavodiiron protein. APX – Ascorbate peroxidase. GST – Glutathione S transferase. PTOX – Plastid/Plastoquinol terminal oxidase.

To additionally understand whether the strong change in the detoxification enzyme abundance was not simply a reason of their absence in LL, we relied on absolute quantification of the enzyme concentration. To this end, intensity-based absolute quantification (iBAQ) (Schwanhäusser et al., 2011) was used (Fig. 1B). Out of the abovementioned enzymes, SOD was the most abundant (top 0.7% percentile compared to all detected proteins), followed by GPX (6.7%) and CBR (7.9%). (Fig. 1B). This further highlights the contribution of CBR and enzymatic antioxidants for photoprotection in HL-grown *C. ohadii* cells.

### HL-grown cells accumulate less oxidatively damaged proteins

To functionally assess the effects of ROS detoxification, the extent of oxidative damage was assessed by quantifying the irreversible oxidation of tryptophan – itself leading to protein degradation (Kato et al., 2023). Remarkably, there were more oxidized peptides from PSII, LHCII, PSI, and LHCI subunits, accumulating in LL-grown cells when compared to HL-grown cells (Fig. 1C), despite the 20-fold lower illumination intensity. This observation underlines the high efficiency of ROS scavenging in HL-grown cells. Taken together, our proteomics analysis suggests that, in addition to the robust downregulation of LHCII and marked accumulation of carotenoid in HL-grown cells (Levin et al., 2021; Levin et al., 2023), SOD and GPX – as well as potentially CBR - play a major role in reducing ROS accumulation and oxidative damage in HL-grown cells.

### High light acclimation results in LHCII loss and reduced PSII functional antenna

To understand the extent of losses in LHCII complexes, thylakoid membranes isolated from *C. ohadii* grown under LL and HL conditions were solubilized and separated by centrifugation on sucrose density gradients. As shown in Fig. 2A, LL-grown cells displayed, in addition to a PSI-LHCI supercomplex, a PSII-LHCII super-complex, and some free LHCII. In contrast, while we observed no changes in PSI-LHCI between cells grown at either light regime, we found no PSII-LHCII super-complex in HL-grown *C. ohadii*. In contrast, only the PSII core complex was visible on the gradient.

**Figure 2.**
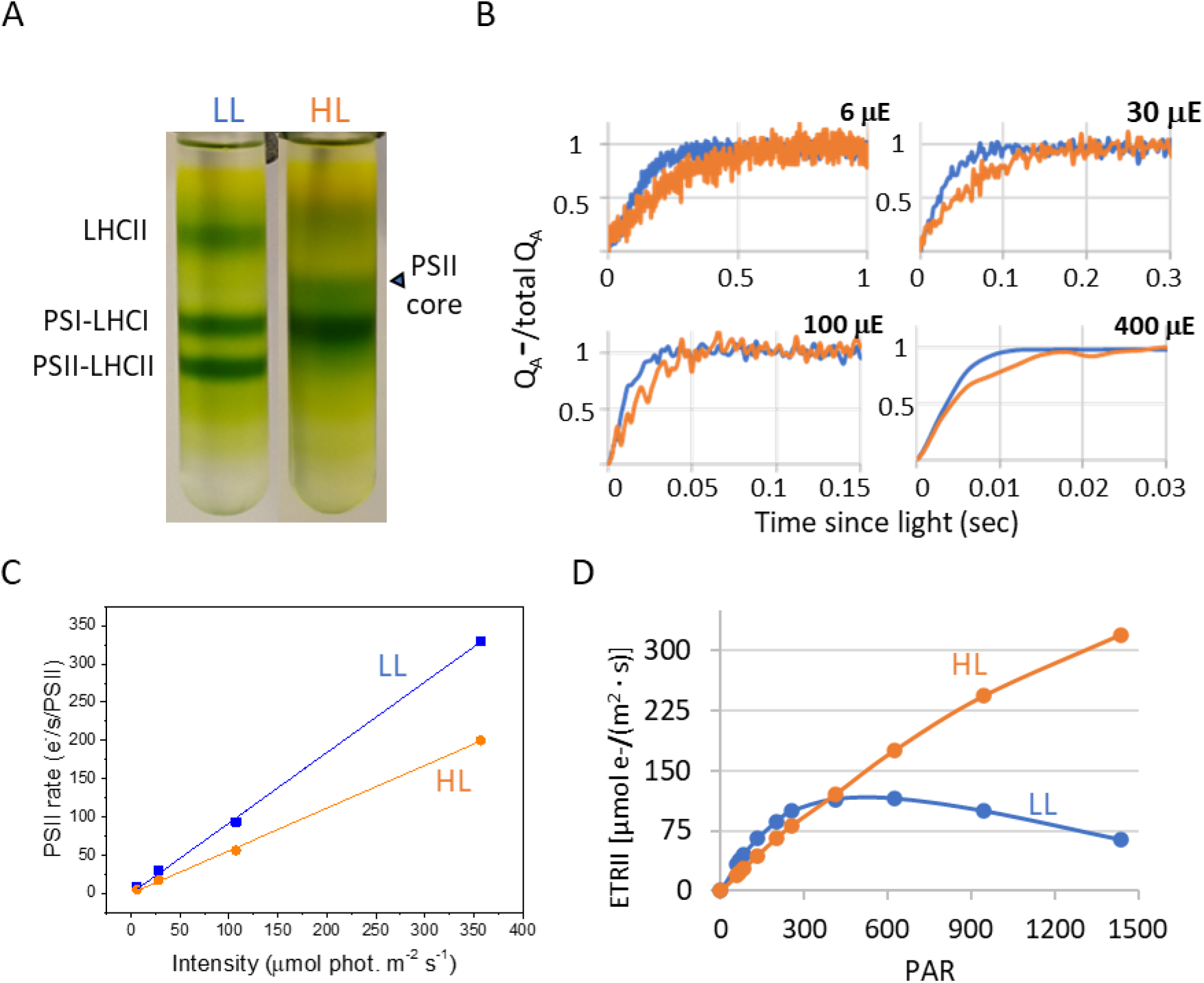
High light-grown *C. ohadii* drastically decreases PSII antenna size. **A)** Thylakoid membranes from low and high-light (LL and HL) grown cells were solubilized with detergents and the photosynthetic complexes were separated by a sucrose density gradient. Samples containing the same amount of chlorophyll (200 µg) were loaded on the gradients. Note the accumulation of a PSII core, stripped from LHCII, in HL-grown cells. **B)** The kinetics of Q_A_ reduction upon exposure to actinic light in the presence of DCMU was measured with a fluorometer and is directly correlated to the PSII antenna size. Y(II) values (Φ_PSII_) were determined under the same experimental conditions, without adding DCMU. Fluorescence data was normalized to maximum (F_M_ = 1) and minimum (F_0_ = 0) values for each sample. The maximal photochemical rate of PSII at each light intensity was calculated by taking the reciprocal of the area delimited by t0, the fluorescence rise curve, and the value of 1 (i.e. F_M_) on the ordinate axis. μE refers to light intensity m^-2^ s^-1^. **C)** Comparison of the slopes of linear regression fits across the measured rate points indicates an approximately 40% decrease in PSII absorption capacity in high-light-grown *C. ohadii*. **D)** Electron transport rates of photosystem II (ETRII) were normalized according to the calculated antenna size and tested under various light intensities. PAR – Photosynthetic active radiation, in μmol photons m^-2^ s^-1^. HL-grown *C. ohadii* ETRII reaches peak values at higher light intensities due to its truncated photosystem II antenna.

To examine how the loss in LHCII and the accumulation of CBR could affect light harvesting in PSII, we performed measurements of its functional antenna size. To this end, we studied the fluorescence rise in the presence of the PSII inhibitor DCMU, which measures the Q_A_ reduction rate upon continuous illumination. The functional antenna size is approximated by calculating the reciprocal of the area above the fluorescence rise curve. A larger area relates to a smaller antenna size, given the slower reduction of Q_A_. As shown in Fig. 2B, the area above the fluorescence rise curves was larger in HL-cells at all actinic light intensities used. The comparison of the slopes of linear regression fits across the measured rate points indicates a 40% difference in PSII absorption capacity between those two conditions (Fig. 2C). This demonstrates that the loss in LHCII induces a strong decrease in the amount of light absorbed by HL-adapted *C. ohadii*. Nonetheless, this is a more limited effect than expected from the massive loss of LHCII observed by the sucrose density gradient assay. These seemingly conflicting observations may be resolved if the CBR protein associated with a large number of carotenoids would act as a PSII antenna that substitutes in part for the loss in LHCII., Alternatively, the LL-grown cells may exhibit already small supercomplex size or accommodate a significant fraction of quenched, free LHCII which do not transfer their excitation energy to the PSII reaction centers.

To understand the effect of a lower PSII antenna size on the PSII electron transfer rate (ETR II) under continuous illumination, the PSII yield (Φ_PSII_) was measured after a 30-second illumination at various light intensities. The yield was then multiplied by the maximal PSII rate at these intensities to provide the ETR II. The results are shown in Fig. 2D and demonstrate that in LL-cells the saturation of PSII ETR is reached at about 400 μmol photons m^-2^ s^-1^, similar to other algae and plants. However, at HL-cells that lack the PSII-antenna, the saturation is reached at much higher light intensities, well above 1500 μmol photons m^-2^ s^-1^, and the ETR amplitude is higher than in LL cells.

### Absence of the q_E_ component of NPQ in Chlorella ohadii

Since the PSII peripheral antenna in green algae and plants is a major site of reversible NPQ, we investigated this photoprotective mechanism in *C. ohadii*. LL- and HL-grown cells were dark-adapted and then subjected to high actinic light intensity (2000 µmol photons m^-2^ s^-1^), after which, their fluorescence behavior were recorded. Strikingly, both LL- and HL-grown cells showed a negligible NPQ response, as indicated by the limited decrease at F_M_’, with little if any NPQ calculated following the equation: NPQ = (F_M_/F_M_’)-1, where F_M_ and F_M_’ are the maximal fluorescence of dark-adapted cells or cells exposed to light, respectively, during a saturating light pulse (Fig. 3A). The decrease in F_M_’ was very limited in *C. ohadii* compared to (HL-grown) *C. reinhardtii* tested under similar light conditions (Fig. 3A). At 2000 µmol photons m^-2^ s^-1^, NPQ reached a value of ∼0.2 (Fig. 3B), in *C. ohadii* compared to ∼2.3 in *Chlamydomonas*, owing to its well established q_E_ NPQ response of high amplitude, facilitated by the proton-sensing capabilities of LhcSR (Fig. 3A). When *C. ohadii* was exposed to even higher light intensities (3000 µmol photons m^-2^ s^-1^), or when grown under strict photoautotrophic conditions, LL-grown cells developed only a limited NPQ response (∼0.7), with similar levels of NPQ in HL-grown cells (∼0.2) (Supplementary Figs. 1B, 1C, and 2)., Thus under all tested conditions, virtually no light-induced q_E_ quenching was detected in *C. ohadii*.

**Figure 3.**
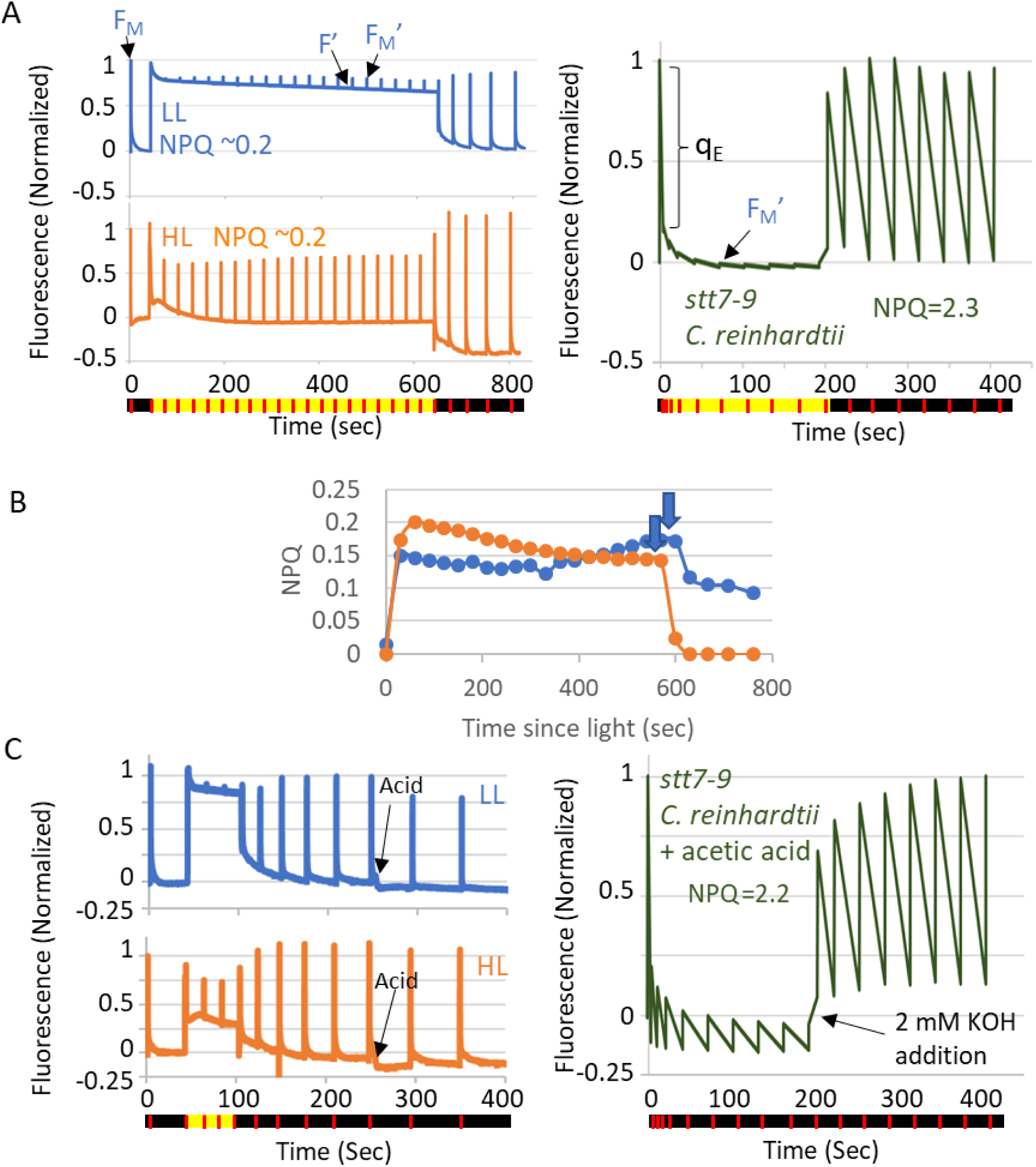
*Chlorella ohadii* exhibits virtually no protective NPQ **A) Left:** Fluorescence traces of dark-adapted LL (top) and HL (bottom) *C. ohadii* grown cells, measured under 2000 μmol photons m^-2^s^-1^. Black and yellow colors at the line below the figure indicate dark and actinic light conditions. The red line indicates the time points when saturating pulses were employed to induce maximum fluorescence (F_M_ and F_M_’). Low NPQ was detected as indicated by the little change in F_M_’ in response to exposure to high light. **Right:** Fluorescence traces of dark-adapted high light grown *stt7-9 C. reinhardtii* cells, exposed to 2000 µmol photons m^-2^ s^-1^. Note the high NPQ levels. **B)** NPQ values were calculated from the fluorescence traces of dark-adapted LL and HL cells as shown in (A). Down-pointing arrows indicate time of turning off of the actinic light. **C)** NPQ is triggered by high light and acidification of the lumen in *C. reinhardtii* but not in *C. ohadii.* **Left:** Fluorescence traces of dark-adapted LL (top) and HL (bottom) *C. ohadii* grown cells, measured under 2000 μmol photons m^-2^s^-1^ or after the acidification of the media to pH 5.5. Black arrows indicate the point of acid addition. **Right:** Fluorescence traces of dark-adapted *stt7-9 C. reinhardtii* cells, exposed to acidic conditions. KOH was added to the cell following 200 sec with no actinic light. The acidification of the medium induced high NPQ levels, mimicking the high light activation of NPQ. This is reversed upon adding KOH, mimicking the light-off action (compare the left panels of **A** and **C**). Fluorescence was normalized between to F_M_ in the dark-adapted cells for all samples.

q_E_ is the major and most rapid component of NPQ which responds to light in seconds and is activated by a pH decrease of the thylakoid lumen. Lumen acidification occurs due to the translocation of protons resulting from high photosynthetic electron flow under HL conditions. These protons bind conserved luminal glutamate residues in LhcSR (algae) and PsbS (plants) and activate q_E_. Even though *C. ohadii* lacks the gene for LhcSR and shows no evidence for PsbS expression (Treves et al., 2016; Levin et al., 2021; Levin et al., 2023), CBRs - which largely accumulate in HL-cells - harbour a conserved glutamate that may function as the proton sensor activating the q_E_ (Levin et al., 2023; Levin and Schuster, 2023). To exclude that there was a limitation to the lumen acidification due to the strong decrease of PSII antenna size, and to verify if CBR can trigger qE at a low lumen pH, we used externally-added acid method, as described in (Tian et al., 2019). The fluorescence traces of LL and HL-grown cells were followed after artificial lumen acidification to pH 5.5. In agreement with the above in vivo analysis of light-induced NPQ, lumen acidification had little effect on the fluorescence signal, indicating the absence of q_E_ activation in both LL- and HL-grown *C. ohadii* cells cultured in either mixotrophic (Fig. 3C) or strictly photoautotrophic (Supplementary Fig. 3) conditions. When the same assay was applied to *C. reinhardtii* cells as a positive control, fluorescence changes, indicative of a strong q_E_ were observed (Fig. 3C). The lack of q_E_ agrees with the absence of the lhcSR gene and the lack of PsbS accumulation in *C. ohadii*, and mirrors the behavior of a *C. reinhardtii* mutant lacking the major NPQ effector, LhcSR3 (Tian et al., 2019), and the psbS mutant of Arabidopsis (Wilson et al., 2024). Importantly, we conclude that despite a conserved glutamate (E131 in *C. ohadii*, E122 in *A. thaliana*) residue on the lumenal side of the predicted CBR structure (Levin et al., 2023; Levin and Schuster, 2023), this protein does not act as a pH-induced q_E_ trigger.

Together, these results indicate that *C. ohadii* does not rely on NPQ mechanisms to survive when grown under intense HL. While LL-grown cells still showed a very limited, slowly developing NPQ effect, possibly due to q_I_ quenching, HL-grown cells were fully lacking in the two types of NPQ.

### HL-grown cells accumulate less damaging ROS than LL-grown cells

To quantify the photoprotective capacity of *C. ohadii* to limit ROS accumulation in the absence of NPQ, we used confocal microscopy coupled with the detection of Singlet Oxygen Sensor Green (SOSG). This dye emits green fluorescence upon interaction with singlet oxygen (^1^O_2_) (Supplementary Fig. 4). LL- and HL-grown cells were incubated with SOSG and exposed to 10 min of high red actinic light, followed by quantification of the accumulated ^1^O_2_. HL-grown, as compared to LL-grown cells, accumulated significantly lower levels of ^1^O_2_ (Figs. 4A-C). To evaluate the contribution of ^1^O_2_ that was generated via photosynthesis-related processes to this difference, we analyzed the SOSG fluorescence that was emitted specifically from within the chloroplast (Supplementary Fig. 4). The analysis of 24 and 54 individual chloroplasts from LL and HL-grown cells, respectively, showed that individual chloroplasts of LL-grown cells accumulated much higher levels of ^1^O_2_ (Fig. 4C), with a 324% increase following 10 min of red-light illumination, while in HL-grown cells, the increase was limited to an average of 175% (Fig. 4C). These results are in line with the increased accumulation of O_2_^-^ in LL-grown as compared with HL-grown algae, while the two cell types accumulated similar amounts of H_2_O_2_ (Fig. 4D) (Levin et al., 2023). Kinetic analyses further confirmed that LL-grown cells accumulated ^1^O_2_ at a significantly higher rate (Fig. 5). Moreover, a significant proportion of chloroplasts of LL-grown cells accumulated 10 times more ^1^O_2_ than those in HL-grown cells (Fig. 4B).

**Figure 4.**
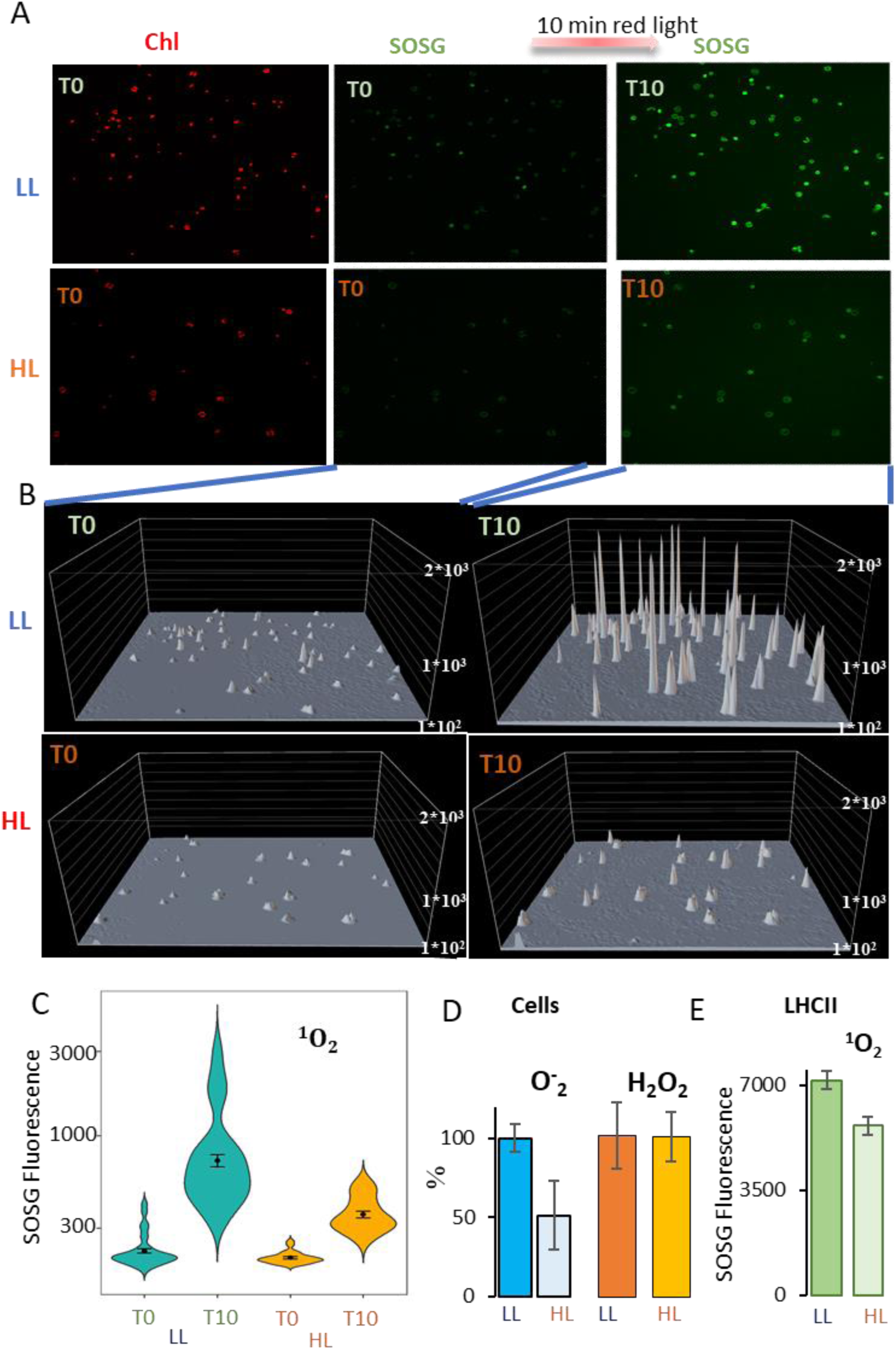
Decreased ^1^O_2_ accumulation in high-light adapted *Chlorella ohadii*. **A)** Confocal microscopy wide-field vision of low and high light (LL and HL)-grown cells. Their intrinsic chlorophyll fluorescence, showed on the left panels, was used to localize the chloroplasts. Note the decreased fluorescence of the HL-grown cells. In the middle and right panels, the fluorescence of the ^1^O_2_ marker SOSG is shown, originating from the same chloroplasts that are shown in the left panels. The SOSG fluorescence has been detected before (T0) and after 10 min (T10) of high red-light treatment. **B)** Quantification of ^1^O_2_ SOSG signal of the chloroplasts shown in A. Only the SOSG fluorescence that overlaps the intrinsic chlorophyll fluorescent marking the chloroplast has been counted for the quantification (see Supplemented Figure 4). **C)** Violin plots of ^1^O_2_ formation following 10 min of illumination with high-intensity red light in 54 and 24 individual chloroplasts of LL and HL-grown cells, respectively. The internal point indicates the mean value and the error bars the standard error (SE). The mean numbers are: LL T0: 223, LL T10: 723, HL T0: 204, HL T10: 358. Note the log scale of the y-axis. The statistical analysis was done as described in the methods section. **D)** Accumulation of O^-^_2_ and H_2_O_2_ in cells after 10 min of strong light illumination. The formation in HL cells was calculated as a percentage of the formation in LL cells (See methods). **E)** ^1^O_2_ accumulation in isolated LHCII complexes after 10 min of strong light treatment was measured by quantifying SOSG fluorescence.

**Figure 5.**
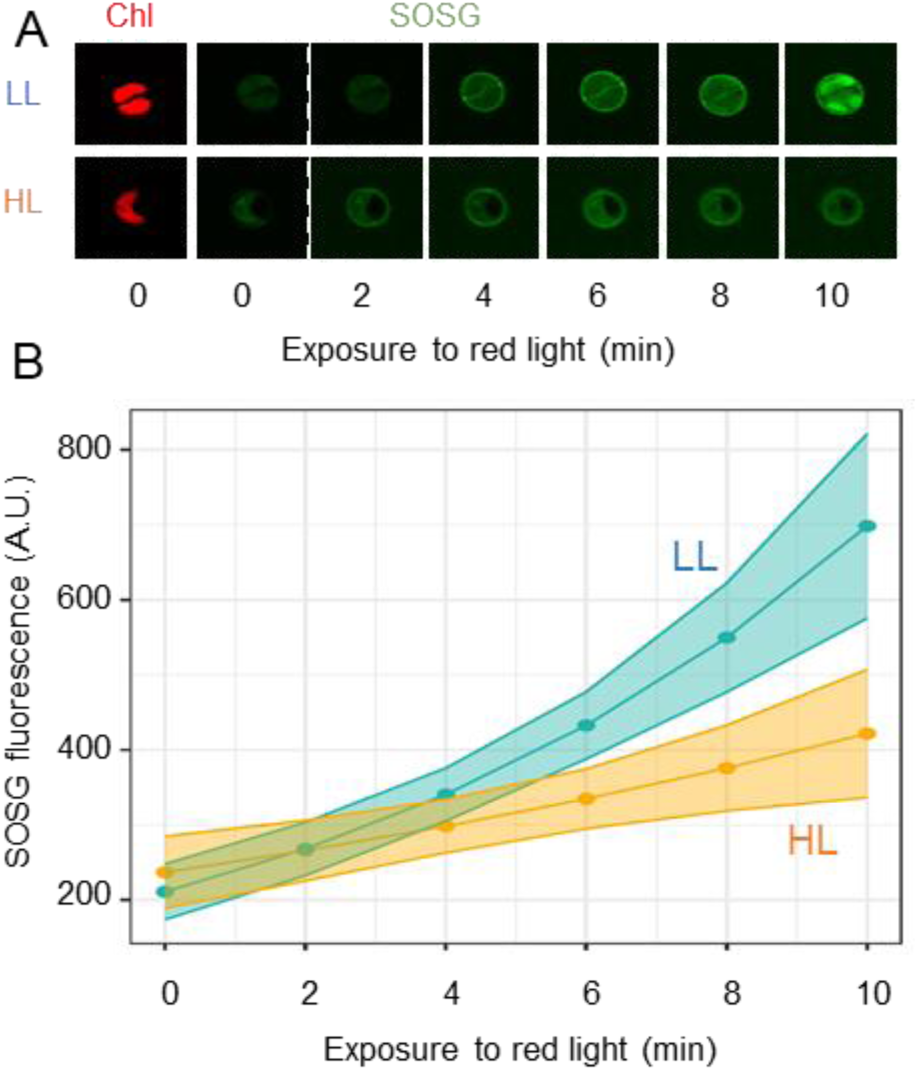
Kinetics of ^1^O_2_ production in *Chlorella ohadii*. **A)** Low and high-light (LL and HL)-grown cells were placed in a medium containing the ^1^O_2_ marker SOSG in the confocal microscope and chlorophyll fluorescence was detected to locate the chloroplasts in the cells (chl). The cells were then illuminated with strong red light for the times indicated at the bottom and the accumulated ^1^O_2_ was detected by the fluorescence intensity of SOSG**. B)** Quantification of the SOSG accumulation in 54 and 24 individual chloroplasts from LL and HL-grown cells, respectively, as shown in panel A. Statistical analysis was performed with a linear mixed model of the log10 transformed response with light as fixed effect and time as both fixed and random effect. Shadowed area represents +/- 2*SE. Mixed model ANOVA showed a significant effect of time (p= 0.0027, table x1) and a significant interaction between light and time (p = 3.215E-06, table x1). Post hoc comparisons between treatments in each time point resulted in significant differences in time points 4,6,8,10 (table x2).

To determine how changes in peripheral antenna between the two light growth conditions were involved in these differences in photosensitivity and ROS production, ^1^O_2_ production was compared between detached LHCII fractions (Fig. 2A) of LL- vs. HL-grown cells at the same chlorophyll concentration. Importantly, detached LHCII fractions from HL-grown cells also contain large amounts of carotenoids and CBR, which may contribute to reduced ^1^O_2_ accumulation via direct ^1^O_2_ scavenging (Levin et al., 2023). The results shown in Fig. 4E demonstrate that the CBR-containing detached LHCII fractions from HL-grown cells indeed accumulated less ^1^O_2_ compared to the same fraction from LL-grown cells. However, the difference was smaller than in whole-cell measurements, suggesting that both the antenna size difference and the differences in the detached antenna properties or the accumulation of CBR and carotenoids had a paramount influence on the rate of ROS synthesis upon photosynthesis.

These observations argue for photoprotective effects mainly originating from the limitation of both light absorption and ^1^O_2_ accumulation by the peripheral antenna.

### Cyclic electron flow is highly activated in HL-grown cells

A decrease in PSII antenna relative to that of PSI creates a favorable environment for CEF around PSI (Fig. 6A) (Tagawa et al. 1963; Joliot and Joliot 2005; Alric 2015; Nawrocki et al. 2019b). This led us assess the kinetics of light-induced P_700_ oxidation to determine whether CEF exhibited a different behavior in LL- vs. HL-grown *C. ohadii*. P_700_ cation absorption at 830 nm serves as a proxy for the activity of CEF when PSII is inactivated by the addition of the herbicide DCMU, or when both CEF and LEF are blocked by DBMIB – a cytochrome *b*_6_*f* inhibitor (Nawrocki et al., 2019a). Drastically different behaviors of P_700_ oxidation were observed between the LL- and HL-grown algae (Fig. 6B). When exposed to the highest tested actinic light intensities in the presence of DCMU, a long lag in P_700_ oxidation was noted when cells were grown in HL, suggesting a high CEF activity (Fig. 6B upper panels). Remarkably, this lag was still observed in presence of methyl viologen, an electron acceptor from PSI, which also decreases the frequency of back reactions while preserving CEF to some extent (Nawrocki et al., 2019a; Joliot et al., 2022). The half-time of PSI oxidation by CEF under these conditions of extremely strong illumination, as determined by the signal obtained following the addition of DCMU, reached 4 seconds, at least 4 times longer than in LL-grown *C. ohadii*, and more than one order of magnitude longer than in *Chlamydomonas* (Nawrocki et al., 2019a). Indeed, the steady state of P_700_ oxidation under continuous illumination at 2000 µmol photons m^-2^ s^-1^– itself a commonly-used proxy for CEF activity (Takahashi et al., 2013; Alric, 2014) - was only around 80%, indicating that even in the absence of PSII activity, a very high rate of CEF can be maintained in *C. ohadii* adapted to HL conditions (Fig. 6B). As expected, this effect was abolished when DBMIB rather than DCMU was added, demonstrating the requirement of electron flow through the cytochrome *b*_6_*f* to maintain CEF. Finally, at lower actinic light intensities in the absence of inhibitors, there was no accumulation of P_700_^+^ due to its efficient reduction by PSII (Fig. 6B). The effect of DCMU was visible in cells grown in LL, while no effect was detected in HL-grown *C. ohadii*, further indicating that a highly efficient CEF can maintain P_700_ mostly reduced. The inability to completely oxidize P_700_ at low light in the presence of DBMIB hints to additional pathways of P_700_^+^ reduction, as substantiated by the relatively rapid reduction of the cation at the end of the illumination period.

**Figure 6.**
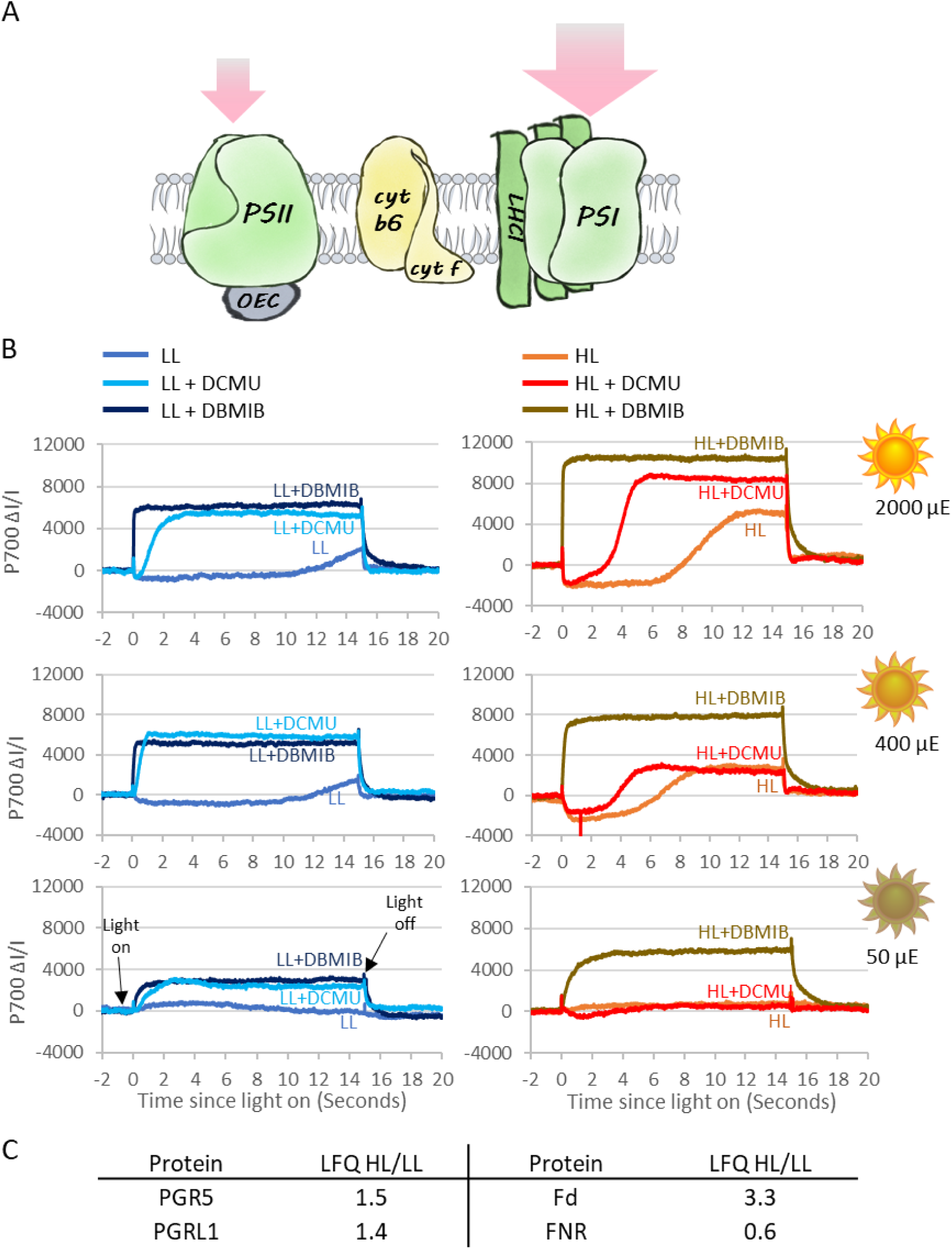
P_700_ redox measurements suggest a strong increase of cyclic electron flow efficiency upon high light acclimation of *C. ohadii*. **A)** In high light (HL)-grown *C. ohadii* the light-harvesting complex II (LHCII) is removed from photosystem II (PSII) while photosystem I (PSI) maintains all or most of its antenna subunits. As a result, PSI is expected to absorb higher quantities of light energy. **B)** P700 oxidation kinetics analysis in dark-adapted cells shows that in the presence of DCMU which blocks linear electron flow (LEF), PSI of HL-grown cells is oxidized at a slower rate compared to that of LL-grown cells in all tested light intensities. When cyclic electron flow (CEF) is blocked by DBMIB, similar PSI oxidation rates are observed in both growth conditions. Note that in LL-grown cells, CEF becomes apparent only by high-light treatment, while in HL-grown cells, CEF is highly activated at all light intensities, and LEF is not activated at very low light intensities. 6 mM methyl viologen was added to all samples. **C)** PGR5, PGRL1, and Fd accumulate in HL-grown cells while FNR accumulates in LL-grown cells, as measured with label-free quantification (LFQ) Liquid chromatography-tandem mass spectrometry (LC-MS/MS).

The stimulation of CEF in HL-grown cells was consistent with mass spectrometry data. According to our proteomics analysis, proton gradient regulation 5 (PGR5) and PRG5-like photosynthetic phenotype 1 (PGRL1), which regulate CEF (Nawrocki et al., 2019b), showed enhanced accumulation in HL-grown cells (1.5-fold for both enzymes) (Fig. 6C). Also, the enhanced accumulation of ferredoxin (Fd) (3.7-fold), together with the decrease in Fd-NADP-reductase (FNR) levels (0.4-fold change), suggested a shift towards CEF rather than to LEF (Fig. 6C).

Together, these results imply that CEF is induced and is a major electron pathway during low light and high light exposure in HL-grown cells, Therefore, aside from the natural requirement to balance the photosynthetic electron flow according to the imbalanced absorption cross-sections of PSII and PSI, the highly active CEF in HL-grown cells likely contributes to extra ATP generation that would be required in these environmental conditions.

## Discussion

### An efficient NPQ-independent photoprotective mechanism underlines *C. ohadii* remarkable resistance to high-light stress

Extremophile species are fascinating forms of living matter, even more so that - their growth environments being exacerbated versions of stressful situations - they may serve as models for the discovery of engineering strategies for the generation of more resistant crops. Photoprotection is thought to be a primary photosynthesis defense mechanism. The various forms of NPQ-controlled photoprotection processes are widely studied, from their molecular mechanism, by numerous molecular and chemical effectors, to their impact on gene expression (for recent reviews, see (Niyogi and Truong, 2013; Murchie and Ruban, 2020; Bassi and Dall’Osto, 2021)). While similar NPQ pathways are widespread in most plants and algae, the current work shows that an extremely light-tolerant microalga, *C. ohadii*, does not require any NPQ to thrive under extreme HL intensities (Fig. 7). When these cells are grown at LL intensities, PSII harvests light both through its core and peripheral antennae and generates ROS, as the membranes contain limited amount of carotenoids (Fig. 7A). However, when these LL-grown cells are exposed to HL intensities before being acclimated to HL conditions, even more ROS are produced, leading to protein oxidation and a high-light acclimation response (Fig. 7B). Indeed, this acclimation is efficient: we observed large differences in the levels of oxidized peptides between the LL and HL cells, suggestive of higher levels of protein damage in LL-grown cells (Fig. 1C). Acclimation to HL conditions leads to a major loss in PSII peripheral antennae while several antioxidant enzymes, CBR, and a large number of carotenoids accumulate. The combined effect of these processes is a marked decrease in the generation and accumulation of ROS, which results in less protein oxidation and turnover (Fig. 7C). Thus, although the elimination of the LHCII in HL-grown *C. ohadii* leads to a potential loss of NPQ sites, it significantly reduces the absorption cross-section of PSII which, in turn, keeps ROS production to a low level under high light. Together with ROS detoxification carried out by the increased accumulation of antioxidants, it drastically reduces the rate of damage that could be caused by the concurrent energy conversion process (Fig. 7).

**Figure 7.**
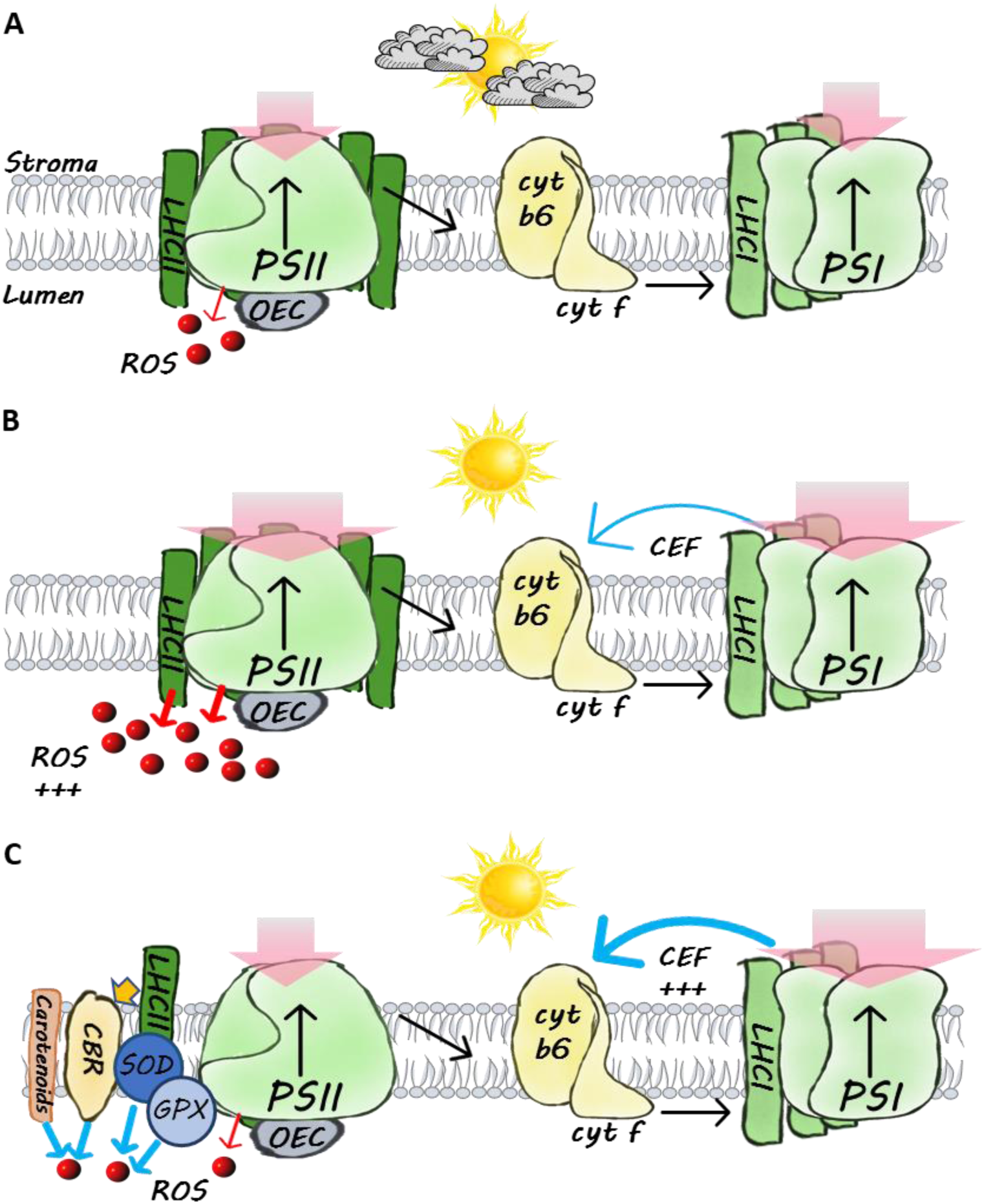
*Chlorella ohadii* does not require NPQ for photoprotection. **A)** Under low light (LL) conditions, linear electron flow is favored and small amounts of reactive oxygen species (ROS) are generated. **B)** Upon a shift to high light (HL) conditions, large amounts of ROS are produced in the photosystem II (PSII) and by its antenna the light-harvesting complex II (LHCII). Cyclic electron flow (CEF) around PSI is relatively low. **C)** To avoid light-induced oxidative damage, HL-grown *C. ohadii* eliminates the LHCII, massively reducing the absorption of excess light energy. Additionally, carotenoids, superoxide dismutase (SOD), glutathione peroxidase (GPX), and possibly the carotenoid biosynthesis-related (CBR) protein and other enzymatic antioxidants act as ROS scavengers. CBR may quench light energy that is absorbed by the detached LHCII, before their degradation. CEF is highly active – at least in the absence of LEF - and may provide extra ATP for metabolic processes.

### Minimizing light absorption may be a more efficient approach to photoprotection than NPQ induction under steady illumination

A drop in LHCII is potentially a risky acclimation strategy for photoprotection upon exposure to high light conditions for at least two reasons. It leads to a loss in most NPQ regulatory processes which are critical to mitigate the effects abrupt changes in light intensities under a fluctuating light regime. In addition, it decreases the light-harvesting capabilities during LL periods. However, *C. ohadii* thrives in the soil crusts of the desert under direct sunlight, where few clouds or vegetation can provide shading. Therefore it is placed in a very stable light environment with extended time periods under constant HL. In these conditions the small core antenna of PSII provides enough light energy to fuel PSII-mediated electron transfer and elimination of LHCII allows a drastic reduction in light-induced ROS production (Fig. 7). It is of note that alongside LHCII elimination, the loss of NPQ in *C. ohadii* also is driven by the absence of the LhcSR gene and PsbS protein (Treves et al., 2016; Levin et al., 2023). Thus, *C. ohadii* provides a remarkable demonstration of what NPQ is not suited for: it does not deliver an appropriate photoprotection against long-lived illumination at very high light intensities, as encountered among extremophiles. NPQ should be regarded as a series of photoprotection responses that evolved with oxygenic photosynthesis exposed to fluctuating light ranging from frequent non-saturating illumination to transient exposures to saturating light as is the case for most crop species and microalgae studied so far. This interpretation is in line with the recent data of photosynthetic response in the frequency domain, with strongest NPQ effect taking place on ∼minutes timescale (Niu et al., 2023).

The extent of the antenna loss, estimated from mass spectrometry data upon solubilization of photosynthetic membranes, looked more extensive that the significant but more limited ∼40% functional decrease in PSII light harvesting capacity during a switch between LL and HL growth (Figs. 1 and 2). This apparent discrepancy may stem from a functional antenna size in LL conditions being smaller than suggested by the amount of LHCII in the sample, in which case a fraction of this peripheral antenna would be disconnected from the reaction centers but aggregated in a quenched state. These quenched LHCII would not contribute to PSII cross-section, and they would barely affect the F_V_/F_M_ in cells, and their loss upon HL acclimation would be consistent with the decrease of chlorophyll content. As an alternative hypothesis, a substitution mechanism for LHCII may take place under HL condition: another light-harvesting complex would contribute to the PSII absorption cross-section, CBRs being the primary candidates given their impressive increased abundance. In the future, time- and spectrally-resolved fluorescence methods could reveal the lifetimes of chlorophyll excited states in each fraction of the antenna in vivo and help decipher which of the two hypotheses is correct.

### Minimizing light absorbance while accumulating antioxidants leads to reduced oxidative damage

In addition to lowering ROS production, the decrease in LHCII content in *C. ohadii* is accompanied by the accumulation of antioxidants such as carotenoids (Levin et al., 2021; Levin et al., 2023) and specialized enzymes, specifically GPX, SOD, and CBR (Fig. 1A). The role of the latter protein remains elusive, but the marked increase in accumulation under HL suggests a key role in *C. ohadii* photoacclimation (Fig. 1A). Despite its structural similarities with PsbS and LhcSR, CBR does not induce q_E_ or other NPQ processes via lumen acidity sensing (Fig. 3B and Supplementary Fig. 3). It was suggested that CBR binds carotenoids (Levy et al., 1993) and, indeed, it was found to co-localize with carotenoids and the detached LHCII fraction of HL-grown *C. ohadii* (Levin et al., 2023). Carotenoids are efficient ROS scavengers (Triantaphylidès and Havaux, 2009), and CBR may contribute to these activities while it is bound to carotenoids. Moreover, the significant accumulation of carotenoids in the thylakoid membrane of HL-grown *C. ohadii*, predominantly zeaxanthin, and lutein, may contribute to this effect (Levin et al., 2023). In addition to being a ROS target, CBR may be an internal UV light screen, expressed by the cells to prevent donor-side damage to PSII.

The overall level of detectable ROS was significantly decreased upon illumination of HL-grown alga. Given the elimination of the PSII antenna and antioxidant accumulation, it is likely that the decline in ROS levels is a product of the smaller PSII absorption cross-section and of the detoxifying effect of the upregulated ROS-scavenging enzymes (Fig. 7). More specifically, the smaller cross-section results in a mild decrease in the amount of absorbed light, which limits ROS synthesis (Santabarbara et al., 1999; Santabarbara et al., 2001). Then, the decreased concentration of nascent ROS due to scavenging activity or using CBR-bound carotenoids as an oxidation target, minimizes the overall accumulation of these deleterious species.

### What is the role of highly active cyclic electron flow in *C. ohadii*?

Finally, CEF around PSI is often thought of as a transient process that occurs before the Calvin-Benson cycle becomes active at the onset of illumination of vascular plants. In addition, because CEF drives acidification of the thylakoid lumen, it contributes to photoprotection of PSII through NPQ activation but also to protecting PSI from photoinhibition since it slows down P_700_ ^+^ re-reduction by the cytochrome *b_6_f* complex (Harris and Boekema, 2018). Last, it has also been implicated at a means to maintain ATP production in anaerobic conditions in microalgae, when mitochondrial respiration is inhibited (for recent reviews on CEF see (Suorsa, 2015; Labs et al., 2016; Nawrocki et al., 2019b)) The finding that CEF is highly active particularly in HL-grown *C. ohadii* calls for a specific role in this alga. Since, little or no lumen acidification-induced NPQ was detected in the alga (Fig. 3B), it shows that this highly active CEF is not used to boost PSII photoprotection by the classic NPQ mechanism. Additionally, P_700_ is primarily maintained in its neutral form even under high light (Fig. 6B)(Caspy et al., 2021). These findings suggest that CEF is not employed to boost lumen acidification – neither for PSII, nor for PSI photoprotection.

It remains that CEF in *C. ohadii* should enhance ATP synthesis in HL conditions. This may be needed to cope with high rates of photoinhibition under desert growth conditions and the heavy energetic cost of PSII repair. Generally speaking, CEF-generated ATP would provide the required energy for keeping with intracellular metabolism in non-growing or slowly-growing conditions, whether for protein and DNA repair, protein translation and energy storage for dark periods.

It is a long-standing technical difficulty in the field of functional photosynthesis to ascertain that transient CEF in the absence of PSII activity – as is the case in our measurements – reflects the situation when LEF is functioning (Fan et al., 2016). The long-lived CEF at the onset of illumination in *C. ohadii* could nonetheless serve as a model for the development of novel methodologies for its quantification, a critical feat for photosynthesis physiology in the coming years.

## Materials and Methods

### Cell culture and acclimation to low or high light conditions

Exponential-phase *Chlorella ohadii* cells were diluted to optical density (OD)_750nm_ 0.2 in a final volume of 400 ml tris-acetate phosphate (TAP) medium and then split: 200 ml were placed under LL (50 µmol photons m^-2^ s^-1^) conditions and 200 ml were placed under HL (2,000 µmol photons m^-2^ s^-1^) conditions, in 1 L Erlenmeyer under shaking (100 rpm), for 24 h, at 28 °C. To prevent self-shading, the cells were maintained below an OD_750nm_ value of 0.8 by diluting them to OD_750nm_ 0.2 every 8 h during the growth period. After 24 h, the LL- and HL-grown cells were harvested for further experimentation. For experiments under strict photoautotrophic conditions, the cells were grown in a tris-phosphate (TP) medium, with no acetate as a carbon source.

Chlamydomonas stt7-9 strain, lacking the kinase activity required for State transitions, was grown in continuous 400 µmol photons m^-2^ s^-1^ condition in MIN media.

### Photosynthetic complexes separation by sucrose density gradient centrifugation

LL- and HL-grown cells were pelleted and washed twice with sucrose (300 mM)-tris-tricine (15 mM each pH 8)-NaCl_2_ (15 mM) (STN) buffer. The cells were broken in a MicroFluidizer (50 psi); unbroken cells were pelleted (15,000 g, 10 min, 4 °C). The supernatant was collected and thylakoid membranes were pelleted in an ultracentrifuge (250,000g, 2 h, 4 °C). Thylakoid membranes were resuspended in 25 mM MES (pH 6.5) to a chlorophyll concentration of 0.2 or 0.4 mg/ml for LL- or HL-grown samples, respectively, and solubilized with 1% *n*-dodecyl-α-D-maltopyranoside (α-DDM) for 15 min, on ice, under dark conditions. Insolubilized material was pelleted (9,500 g, 10 min, 4 °C) and the supernatant was loaded on a sucrose density gradient (20 mM MES pH 6.5, 0.02% α-DDM, 4%-45% sucrose). Individual complexes were separated by ultracentrifugation (100,000 g, 20 h, 4 °C)

### Functional analyses

LL- and HL-grown cells were pelleted and resuspended in fresh medium to the equivalent of 10 μg/ml chlorophyll for PSII chlorophyll fluorescence analysis, or 50 μg/ml for P_700_ (PSI) absorption analysis. The cells were dark-adapted for at least 30 min with air bubbling, then loaded to a quartz cuvette and analyzed with a Dual-PAM-100 fluorometer (Walz, Germany). Red actinic light was used in all measurements.

#### PSII electron transport rate saturation curve

After exposure of LL- and HL-grown cells to different light intensities for 30 sec, the PSII operating efficiency (Φ_PSII_) was measured using the following equation: Φ_PSII_ = (F_M_’-F’)/F_M_’. PSII ETR was calculated following the equation: ETR(II) = σ_II_ · Φ_PSII_, where σ_II_ is the maximal, light-limited rate of PSII at each light intensity (“PSII antenna size”). The latter is the reciprocal of the area above fluorescence-DCMU curve, delimited by F_M_ value normalized to 1 and by the t_0_.

#### NPQ analysis

The fluorescence traces of LL- and HL-grown cells were monitored upon exposure to various light intensities (50, 400, 2000, and 3000 µmol photons m^-2^ s^-1^) for 10 min, followed by 3 min of darkness for recovery. When indicated, acetic acid (1 mM final) was added directly to the cuvette to induce q_E_ (Tian et al., 2019). NPQ was calculated following the equation: NPQ = (F_M_/F_M_’)-1 in both *C. ohadii* and *C. reinhardtii*.

#### P_700_ redox measurements

P_700_ oxidation kinetics were monitored in LL- and HL-grown cells upon illumination with various light intensities: 50, 400, or 2000 μmol photons m^−2^ s^−1^, in the presence or absence of the photosynthetic electron transfer inhibitors 3-(3,4-dichlorophenyl)-1,1-dimethylurea (100 μM, DCMU) or 2,5-dibromo-3-methyl-6-isopropylbenzoquinone (100 μM, DBMIB). Methyl viologen (6 mM) was added to all samples to prevent charge recombination in PSI.

### ROS accumulation assay

#### Live-cell ^1^O_2_ quantification with singlet oxygen sensor green

LL- and HL-grown *C. ohadii* cells were pelleted and resuspended to OD_750nm_ 0.2 in TAP medium containing 50μM singlet oxygen sensor green (SOSG). Subsequently, the cells were incubated for 30 min under LL conditions, at 28°C, and pelleted and resuspended in fresh TAP medium. To visualize the formation of ^1^O_2_, the cells were loaded to a spinning disk confocal microscope (Nikon Eclipse Ti2, Confocal scanner unit Yokogawa CSU-W1, Camera Photometrics PrimS-BSI). Light stimulation was achieved using a red LED integrated into the microscope, providing an intensity of 3.9 mW (660 nm) for a duration of 1-10 min. The SOSG was excited at 488 nm, and the fluorescence emission was recorded at 530 nm. Chlorophyll fluorescence was recorded with excitation at 430 nm and emission at 680 nm to identify the chloroplasts. IMARIS software (Oxford Instruments) was employed to reconstruct the chloroplast structure based on the chlorophyll fluorescence. Subsequently, only SOSG fluorescence that corresponded to the reconstructed chloroplast was measured, ensuring a specific and accurate assessment of the effects of light stress on the observed photosynthetically derived ^1^O_2_ formation. The graphs were prepared with the R environment for statistical computing and graphics (https://www.R-project.org/) using the ggplot2 package (Wickham, 2016). The linear mixed model was calculated with the lme4 package (Bates et al., 2015). SEs and post-hoc analysis were calculated using the emmeans package (https://CRAN.R-project.org/package=emmeans). Mixed-model ANOVA was used to determine statistical significance (see Fig. 5B)

#### ^1^O_2_ accumulation in isolated LHCII complexes

Isolated LHCII from LL-grown cells and isolated LHCII together with CBR from HL-grown cells were resuspended in STNM buffer (50 mM Tricine-KOH pH 7.6, 15mM NaCl_2_, 300 mM sucrose, and 5mM MgCl_2_) with SOSG (10 mM) to a final chlorophyll concentration of 0.028 μg/μl. The samples were loaded onto a black 96-well plate and SOSG fluorescence was measured before and after 10 min of light illumination in a fluorescence plate reader (ClarioStar 500, BMG Labtech, Ortenberg, Germany) with excitation at 488 nm and an emission range of 530-600nm. The samples were placed on ice during the illumination period to avoid protein denaturation. STNM buffer with SOSG was used as a blank to measure and exclude the background SOSG fluorescence. Three biological repeats were tested for each treatment condition.

#### O_2_^-^ and H_2_O_2_ accumulation in low-light- and high-light-grown C. ohadii cells

Nitro-blue tetrazolium (NBT), which upon reduction creates blueish–greyish insoluble precipitates, and 3,3′-diaminobenzidine (DAB), which upon oxidation by H_2_O_2_ becomes dark brown, were used to measure O_2_^-^ and H_2_O_2_ accumulation in LL- and HL-grown *C. ohadii* cells. The cell OD_750nm_ was adjusted to 0.7. One millilitre of each sample was subjected to a 10 min incubation under HL conditions, with the addition of either 5 mM NBT or 5 mM DAB. Thereafter, chlorophyll was extracted with 100% methanol, at 40°C. Cells were then pelleted (3000 g, 5 min), and resuspended in 200 μl methanol (100%). The resulting cell suspension was applied as spots onto the Whatman 3 paper filter. ROS were quantified with ImageJ, based on the colour intensity.

### Mass spectrometry whole-cell sample preparation and proteolysis

Three biological repeats of LL- and HL-grown *C. ohadii* cells were harvested, washed twice with STN, and broken under 50 psi with a MicroFluidizer. Samples were suspended in 8.5 M urea, 100 mM ammonium bicarbonate, and 10 mM DTT. Protein content was estimated using Bradford readings. The samples were reduced (60 °C, 30 min), modified with 35.2 mM iodoacetamide in 100 mM ammonium bicarbonate (room temperature for 30 min in the dark), and digested in 1.5 M urea and 17.6 mM ammonium bicarbonate with modified trypsin (Promega), overnight, at 37 °C, at a 1:50 (M/M) enzyme-to-substrate ratio. A second digestion with trypsin was performed for 4 h, at 37 °C, at a 1:100 (M/M) enzyme-to-substrate ratio. The tryptic peptides were desalted using Oasis HLB 96-well µElution Plate (Waters), dried, and re-suspended in 0.1% formic acid.

### Mass spectrometry analysis

The resulting peptides were analysed by LC-MS/MS using a Q Exactive HFX mass spectrometer (Thermo) fitted with an HPLC capillary (Ultimate 3000, Thermo Scientific). The peptides were loaded in solvent A (0.1% formic acid in water) on a homemade capillary column (30 cm, 75-micron ID) packed with Reprosil C18-Aqua (Dr. Maisch GmbH, Germany). The peptide mixture was resolved with a 5-28% linear gradient of solvent B (99.99% acetonitrile with 0.1% formic acid) for 180 min, followed by a gradient of 28-95% for 15 min, and then 15 minutes at 95% acetonitrile with 0.1% formic acid in water at flow rates of 0.15 μl/min. Mass spectrometry was performed in a positive mode (m/z 350-1200, resolution 120,000 for MS1 and 15,000 for MS2), using repetitively full MS scan followed by high collision dissociation (HCD, at 27 normalized collision energy) of the 30 most-dominant ions (>1 charges) selected from the first MS scan. The AGC settings were 3x10^6^ for the full MS and 1x10^5^ for the MS/MS scans. A dynamic exclusion list was enabled with an exclusion duration of 20 s.

### Mass spectrometry data analysis

Raw data was processed with MaxQuant version 2.5.2 (Tyanova et al., 2016) for protein accumulation analysis and MSfragger version 4.0 (Kong et al., 2017) via FragPipe version 21.1 (https://fragpipe.nesvilab.org/) for detection of tryptophan oxidized peptides. In the absence of a fully annotated proteome for *C. ohadii*, we utilized a combined dataset of protein sequences in our identification process. This dataset includes all available Chlorella protein sequences (Tax ID = 3701) downloaded from UniProt as of July 2019, comprising a total of 20973 sequences. Additionally, we incorporated all available *C. ohadii* protein sequences (11470) from NCBI, as downloaded on October 2023. All searches included enzymatic cleavage with trypsin (cleavage after K/R but not P) and two missed cleavages and a clip of N-term methionine unless specified otherwise. Cysteine carbamidomethyl (+57.02146) was set as a fixed modification, and methionine oxidation (+15.994915) and protein N-terminal acetylation were set as variable modifications.

MaxQuant searches were performed using tryptic digestion mode with a minimal peptide length of 7 amino acids. Search criteria included up to 2 missed cleavages, precursor and fragments tolerance of 20 ppm, oxidation of methionine, and protein N-terminal acetylation set as variable modifications. Default settings were used for all other parameters. Candidates were filtered to obtain an FDR of 1% at the peptide and protein levels. For quantification, the match between runs (MBR) module of MaxQuant was used with the Label-Free Quantification (LFQ) normalization method enabled.

Identification and quantification of tryptophan oxidation was performed using FragPipe’s ‘LFQ-MBR’ workflow. The workflow default settings were used with the additions of tryptophan oxidation (+15.994915) and deoxidation (+31.989829) as variable modifications for identifying tryptophan oxidation.

Further analysis was performed using the Perseus software v.2.0.11 (Tyanova et al., 2016). Label-free quantification (LFQ) intensity values were used to calculate the relative abundance of total proteins and tryptophan oxidized peptides in LL and HL whole-cell samples. The results were filtered to exclude contaminants, reverse, and only identified by site proteins. Label-free quantification values were transformed (Log2) and filtered to include only proteins that were detected in three repeats of at least one group: LL or HL samples. Missing values were then added from a normal distribution (https://cox-labs.github.io/coxdocs/replace-missingfromgaussian.html). A two-sample t-test was used to determine the statistical significance of differences between groups, applying a permutation-based false discovery rate (FDR) of 0.05 and an S0 parameter of 0.1. Intensity-based absolute quantification (iBAQ) (Schwanhäusser et al., 2011) was used to determine the abundance of a given protein in HL-grown cells. A protein with a higher iBAQ value is more abundant compared to a protein with a low iBAQ value. The proteins were listed according to their average iBAQ values (high to low) in HL-grown cells. The percentile of the protein was determined by dividing its number (position) on the list, by the total number of proteins, and multiplying by 100. For example, if protein X is 40^th^ on the list, and a total of 4000 proteins were detected in the analysis, then protein X is in the 1^st^ percentile, i.e. in the top 1% most abundant proteins in the cell.

## Data availability

The mass spectrometry proteomics RAW data have been deposited to the ProteomeXchange Consortium via the PRIDE (Perez-Riverol et al., 2022) partner repository with the dataset identifier PXD053918

## Acknowledgments

We thank Dr. Nitzan Dahan from the Technion LS&E microscopy unit for his assistance and guidance. We thank the Smoler Proteomics Center at the Technion for performing the mass spectrometry experiments. We thank Nir Keren and Iftach Yacobi for their helpful advice and discussions.

## Author contributions

GL, GS, FAW, and WN planned the research and wrote this manuscript. GL and MS performed the experiments and data analysis unless stated otherwise. OK and RH assisted with proteomics data analysis. GH helped analyze the singlet oxygen accumulation data and performed statistical analyses. OL assisted with NPQ measurements.

## Dedication

This paper is dedicated to the late Professor Itzhak Ohad, an outstanding and beloved scientist and mentor, who made a substantial contribution to our understanding of photoinhibition and paved the way to the isolation of *C. ohadii*.

---

**Supplementary Figure 1.**
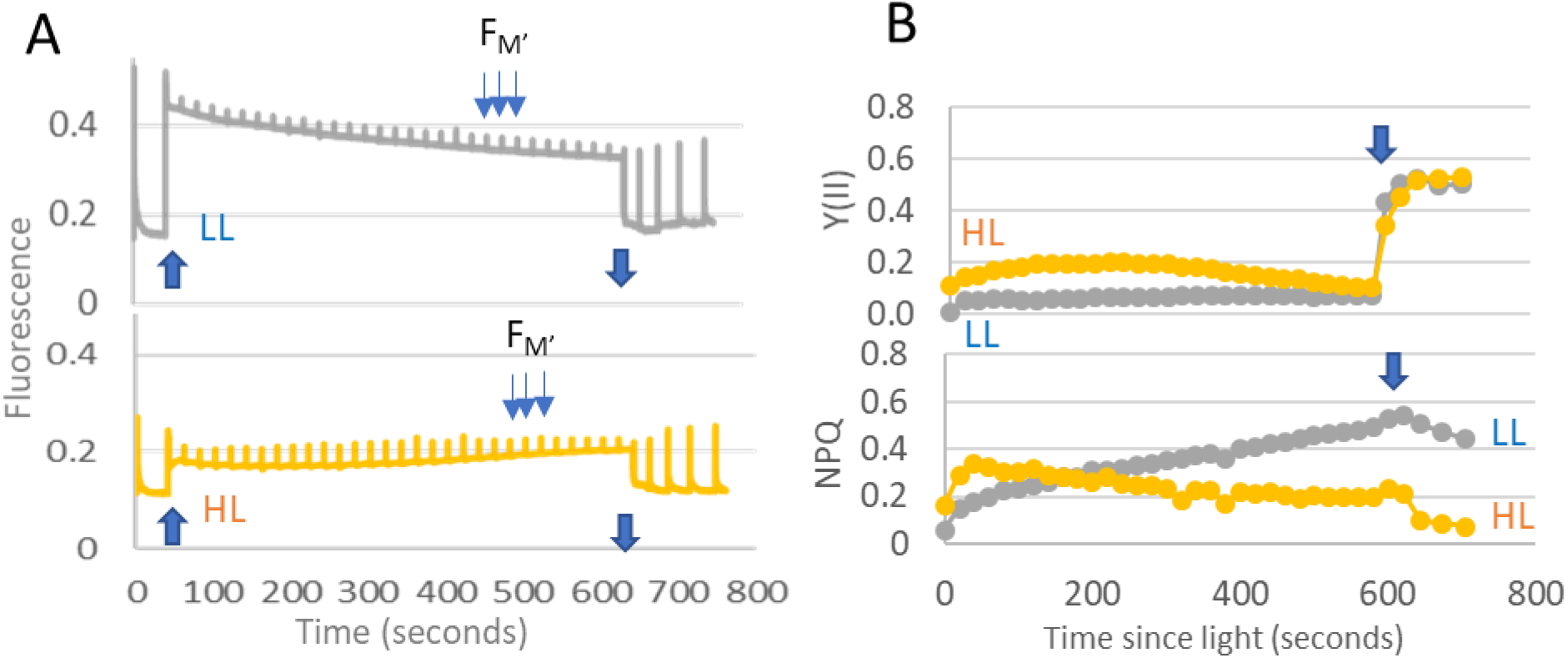
*Chlorella ohadii* lacks NPQ under strict photoautotrophic conditions. **A)** Fluorescence traces of dark-adapted LL (top) and HL (bottom) cells grown in strict photoautotrophic conditions, measured under 2000 μmol photons m^-2^s^-1^. Negligible NPQ was detected as indicated by the little change in maximal fluorescence (F_M_) in LL and no change in HL in response to exposure to the pulses of high light. **B)** NPQ and Y(II) values were calculated from fluorescence signals as shown in panel *A*.

**Supplementary Figure 2.**
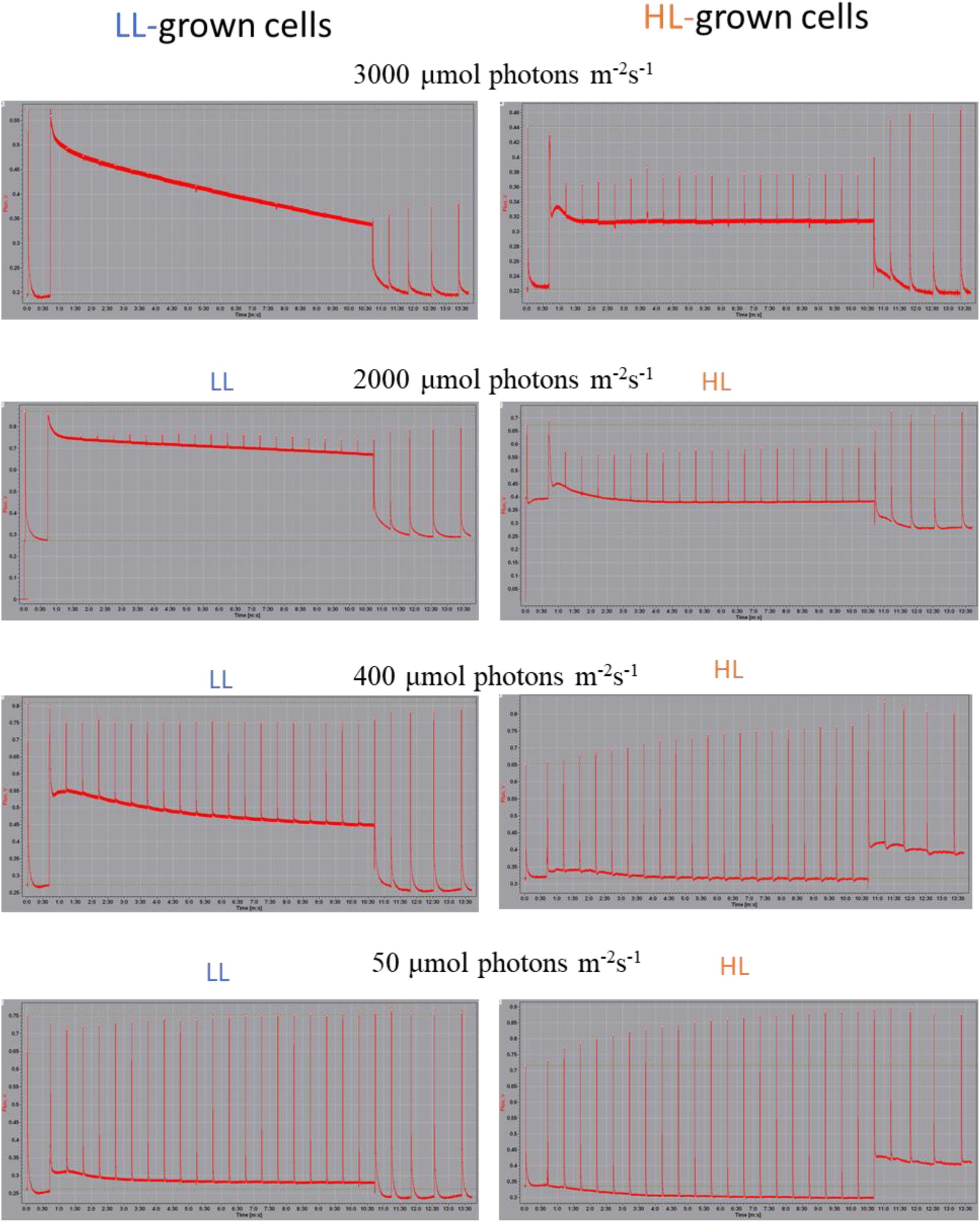
*Chlorella ohadii* lacks NPQ under low or high light intensities. Fluorescence traces of dark-adapted low and high light (LL, left, and HL, right)-grown cells. The fluorescence measurement was carried out under different light intensities, as indicated in the Methods section. Low NPQ was detected as indicated by the little change in F_M_’ in response to exposure to low or high light.

**Supplementary Figure 3.**
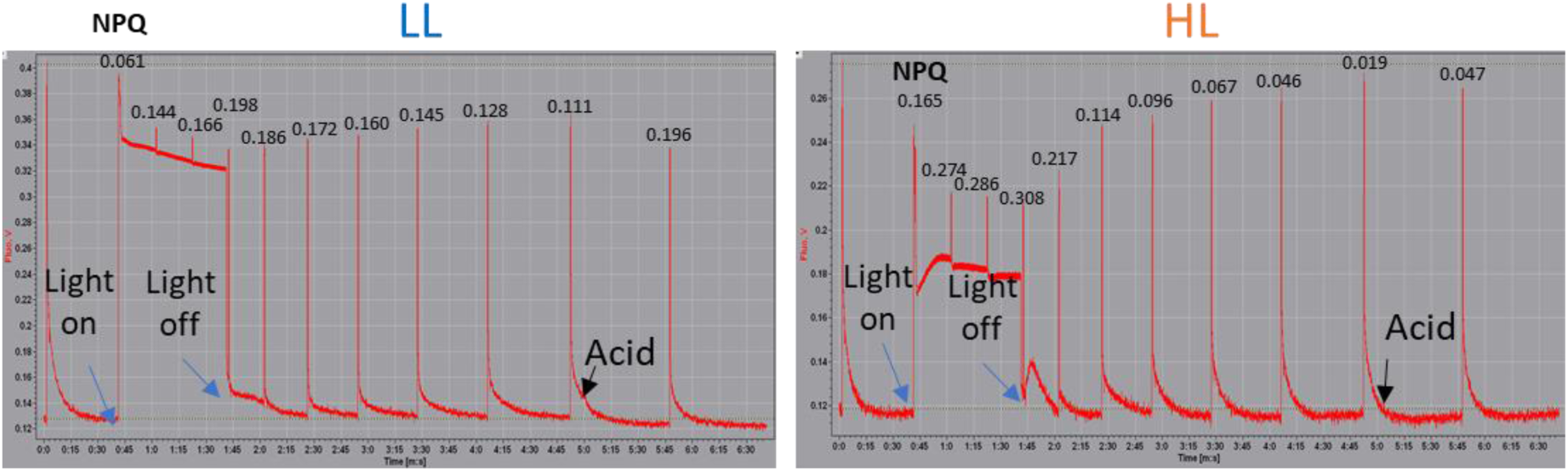
NPQ is not induced by high light or acidifying conditions in *Chlorella ohadii* grown under strict photoautotrophic conditions. Fluorescence traces of dark-adapted low and high light (LL, left, and HL, right)-grown cells. First under high light exposure (2000 μmol photons m^-2^s^-1^), and then quenching was detected in response to the acidification of the media to pH 5.5 with acetic acid. Blue arrows indicate turning the light on/off. Black arrows indicate the point of acid addition. The numbers above each FM signal indicate the NPQ values which are very small compared to *C. reinhardtii* where they routinely attain 2.2-2.5 (see Figure 3).

**Supplementary Figure 4.**
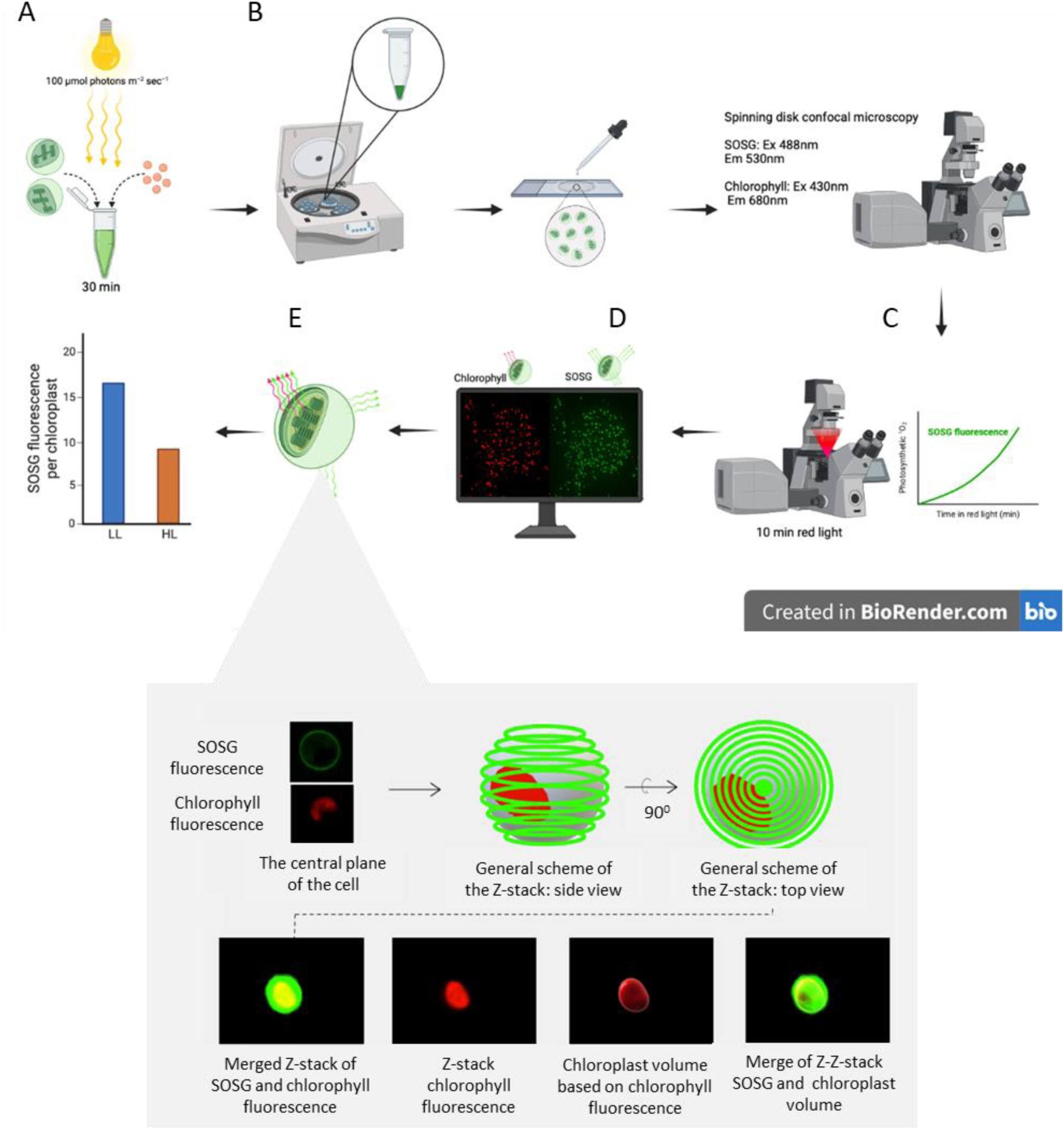
^1^O_2_ detection with singlet oxygen sensor green (SOSG) probe workflow. **A)** SOSG was introduced to LL and HL cells by incubation for 30 min at the light intensity of 100 μmol photons m^-2^ sec^-1^, with gentle shaking. **B)** The cells were pelleted and resuspended in a fresh TAP medium and loaded onto a microscope slide and applied to a spinning disk confocal microscope with a Z stacking of 8 μm around the cell central plain. **C)** Induction of high light stress was initiated within the microscope using red LED illumination. This wavelength selection (∼600 nm) was chosen to specifically activate photosynthesis. **D)** SOSG fluorescence (excitation at 488 nm/emission at 530 nm) was recorded. To locate the chloroplast, chlorophyll fluorescence (excitation at 430 nm/emission at 680 nm) was concurrently recorded. **E)** Chlorophyll fluorescence was detected in different focused plains before and after the red high light treatment and a 3D model of the fluorescence was created by Z-stacking in the IMARIS program. Due to the limited permeability of SOSG, green fluorescence appeared on the cell membrane throughout the experiments. To subtract the access fluorescence that was not contributed by the photosynthetic-driven ^1^O_2_ formation, a chloroplast volume was constructed from the chlorophyll fluorescence. **F)** SOSG fluorescence was only calculated from the chloroplast volume. This approach ensures the isolation of photosynthetic singlet oxygen formation, offering a comprehensive insight into the dynamics of singlet oxygen accumulation within the cellular context.

